# Long-term multichannel recordings in *Drosophila* flies reveal distinct responses to deviant visual stimuli during sleep compared to wake

**DOI:** 10.1101/2025.01.09.632265

**Authors:** Matthew N. Van De Poll, Bruno van Swinderen

## Abstract

During sleep, behavioral responsiveness to external stimuli is decreased. This classical definition of sleep has been applied effectively across the animal kingdom to identify this common behavioral state in a growing list of creatures, from mammals to invertebrates. Yet it remains unclear whether decreased behavioral responsiveness during sleep is necessarily associated with decreased responsiveness in brain activity, especially in relative latecomers to the sleep field, such as insects. Here, we perform long-term multichannel electrophysiology in tethered *Drosophila melanogaster* flies exposed continuously to flashing visual stimuli. Flies were still able to sleep under these visual stimulation conditions, as determined by traditional immobility duration criteria for the field. Interestingly, local field potentials (LFPs) recorded in a transect through the fly brain failed to show any difference in amplitude during sleep compared to wake when the visual stimuli were invariant. In contrast, LFP responses were lower when visual stimuli were variable and of lower probability, especially in the central brain. Central brain responses to ‘deviant’ stimuli were lowest during the deepest stage of sleep, characterized by more regular proboscis extensions. This shows that the sleeping fly brain processes low-probability visual stimuli in a different way than more repeated stimuli and presents *Drosophila* as a model system for studying the potential role of sleep in regulating predictive processing.

**SUMMARY STATEMENT:** The sleeping fly brain still responds to visual stimuli. However, response amplitudes depend on stimulus history and probability, as well as recording location.

## INTRODUCTION

The key criterion identifying sleep in an animal is decreased responsiveness compared to wake (Campbell and Tobler, 1984; Kales and Rechtschaffen, 1968). Typically, this is investigated in by measuring behavioral responsiveness to probing stimuli, such as mechanical vibrations, loud noises, or light flashes. Responsiveness during sleep can also be tracked in brain activity recordings, where event related potentials (ERPs) in response to applied stimuli are typically lower or even abolished during sleep compared to wake. Even while peripheral sensory processing continues, as in the case of auditory stimuli in many animals, sensory perception can be attenuated during sleep. For example, brain responses to certain auditory stimuli are decreased during sleep in human EEG recordings, revealing impaired predictive processing of unexpected sounds during specific sleep stages (Andrillon and Kouider, 2020; Strauss et al., 2015). This suggests that a key feature of the sleeping brain is a decreased ability to register changes in the external environment. Whether this is true in other animals that sleep is unclear. A recent study in Drosophila melanogaster found that sleeping flies can process and respond to different odors, with ethologically-relevant odors more likely to wake flies up (French et al., 2021). This suggests a level of sensory processing in the sleeping fly brain.

It is well established that flies sleep (Shafer and Keene, 2021), and work over the past decades shows that, like other animals, fly sleep is also characterized by distinct stages (Jagannathan et al., 2024; Tainton-Heap et al., 2021; van Alphen et al., 2021, 2013; Yap et al., 2017). These stages appear to comprise of an ‘active’ phase when the fly brain seems as active as during wakefulness (but behaviorally the flies are quiescent and arousal is low), and a ‘quiet’ phase when brain activity is decreased (Anthoney et al., 2023; Tainton-Heap et al., 2021) and when flies display characteristic microbehaviors such as rhythmic proboscis extensions (Jagannathan et al., 2024). Evidence for altered brain activity during sleep in flies was originally drawn from single-channel local field potential (LFP) recordings, which allowed for long-term recordings in behaving flies (Nitz et al., 2002). In a recent study, we achieved long-term recordings in behaving (and sleeping) flies implanted with a 16-channel silicone probe, which provided insight into how different regions of the fly brain behaved across changes in behavioral state (Jagannathan et al., 2024). These experiments confirmed distinct sleep stages in flies, but surprisingly showed that the entire fly brain sleeps, including the outer optic lobes. How visual stimuli might be processed from the optic lobes to the central brain during sleep is unknown.

Unlike humans and many other animals, flies and other insects cannot shut their eyes when they sleep. Yet, visual responsiveness is suppressed in sleeping insects, as shown in behavioral studies (Campbell and Tobler, 1984) as well as in electrophysiological recordings (Kaiser and Steiner-Kaiser, 1983; van Swinderen and Greenspan, 2003). This suggests a central arousal mechanism designed to block certain visual stimuli during sleep, although it is not known how this mechanism might work or what parts of the insect brain might be involved. In the present study, we adapted our long-term multichannel recording preparation for Drosophila flies exposed to repetitive visual stimuli, to investigate if visual responsiveness across the fly brain changed during sleep. We first examined the effect of steady-state visually evoked potentials (SSVEPs) (Norcia et al., 2015), as done for previous work in awake (Paulk et al., 2013) and anesthetized (Cohen et al., 2016) flies. We then controlled the probability of different visual stimuli to investigate whether visual predictive processing was altered during spontaneous sleep compared to wake. Finally, investigated whether visual processing might be different across the fly brain during distinct stages of sleep.

## MATERIALS AND METHODS

### Animals

To maintain consistency, all animals used in experiments were from a cross of a 23E10-GAL4 driver line (cs; P{y[+t7.7] w[+mC]=GMR23E10-GAL4}attP2) with a UAS-Chrimson line (w[1118]; P{y[+t7.7] w[+mC]=20XUAS-CsChrimson.mCherry}su(Hw)attP5). Animals were raised at 25 degrees on standard cornmeal medium. Female F1 flies were collected between 3 and 9 days post-eclosion (dpe) and used for experiments.

### Tethering

Tethering was performed as described previously (Van De Poll and van Swinderen, 2023). In brief, animals were cold-anesthetized at 2 degrees Celsius and glued dorsally to a short tungsten rod. A silver ground wire was also inserted gently into the thorax. After tethering the animals were placed atop an air-suspended ball.

### Experimental setup

While walking on an air-supported ball, animals were able to view a custom-built 8x8 matrix of interleaved blue and green LEDs, positioned to the front-left and level with the fly. The power of both green and blue LEDs was calibrated to be 0.005mW/mm^2^ at the eye with a photometer.

A 16 channel multielectrode (NeuroNexus A1x16-3mm 50-177) was inserted into the ipsilateral eye to the LED panel, feeding through a preamplifier (Tucker-Davis Technologies RA16PA) into a data acquisition base station (Tucker-Davis Technologies RZ5).

During experiments a camera placed lateral to the fly recorded behavioral activity, including movements of the legs and abdomen, as well as the proboscis. A custom Python script acquired the video data and saved the images as well as CSV timestamps for each frame.

Stimulus paradigms were coded on a computer running Windows 10 (Dell Precision 7910), and then sent to the base station, which simultaneously controlled the transmission of the signal to the power circuit controlling the LEDs (custom-built) and recorded the sent signal via a loopback.

### Stimuli

Stimuli consisted of full-field flashes of the LED panel at a fixed 10Hz interval, with a 50% duty cycle. Each experiment was comprised of a number of ‘trials’, where each trial was defined as a 20s period of stimuli (200 stimulus events). The stimulus signal was a square wave, such that the LEDs moved between the On and Off states with no ramping.

The arrangement of stimulus events within each trial could be one of three combinations, depending on each experiment: “Carrier only” trials consisted of a unitary color (e.g. All green), at the F1 frequency of 10Hz. “Phasic deviant” trials contained a deviation in color (e.g. green deviant amongst blue carriers) at a rate of 2Hz, such that every fifth stimulus event was a deviant. “Jittering deviant” trials were similar, except that the position of the deviants in the sequence was shifted according to a normal distribution (i.e. most likely to be 5^th^ element, less likely to be 4^th^ or 6^th^ element, etc).

### Experiment durations

For initial experiments, the duration was around 1h per fly (72 trials). In the later experiments incorporating sleep analysis, the experiments ran for a portion of the day and a full night cycle, typically around 16h (2450 trials). Flies would slowly perish in this setup, and so typically only the first 6h of recordings were used, based on qualitative observation of signal quality over time for the compiled dataset.

### Data

LFP data was acquired from the inserted multichannel electrode at 24kHz, downsampled after experiments to either 1000Hz or 200Hz for 1h and 16h recordings respectively. In addition to the 16 LFP channels, this data contained time-synchronized signals for the stimulus identity, as well as stimulus color and trial state.

Timestamps were used to synchronize the LFP and behavioral video recordings, to allow for separation of LFP data by activity state and other factors.

### Statistics

All data analysis was performed with custom MATLAB (R2020a) scripts. LFP data was downsampled as mentioned and then line noise was removed at 50Hz multiples. Most data analysis was performed on the SSVEP evoked by each stimulus event, with the primary metric being the delta amplitude between the highest and lowest points of the SSVEP, following the stimulus onset. Normalization was performed within channel by calculating the mean amplitude of all stimulus events for each color (e.g. All green stimuli, regardless of identity) and expressing each individual event as a proportion of that value.

To allow us to compare across conditions during comparisons we calculated the average across groups (within flies) and then subtracted this from individuals, thus yielding a comparison-specific, normalized value, referred to herein as “corrected delta”. For example, in comparing green deviants against carriers, the mean of the deviant and carrier response for each fly would be calculated and then subtracted from the two groups, thus expressing any change as a relative value.

## RESULTS

Recording visual responses across the fly brain during sleep and wake Tethered female Drosophila flies (3-7 days old) were able to walk for hours at a time on an air-supported ball while being presented with flashing visual stimuli (Figure 1A). During this time, local field potential (LFP) activity was recorded from 16 different locations across their brain, spanning from the left retina to the central brain (Figure 1B, top). In the absence of visual stimulation, LFP recordings across the fly brain reveal a characteristic profile, with synchronized endogenous activity across the central brain and optic lobe (Figure 1B, bottom left). As shown in previous studies using a similar multichannel preparation (Paulk et al., 2013), the fly brain responds to periodically flashing visual stimuli with synchronized evoked potentials (Figure 1B, bottom right). A polarity reversal is evident in the vicinity of the lobula (around electrodes 11-14), and this electrophysiological feature was used to ensure reproducible electrode insertions across flies, and to serve as a reference channel (see Methods). As in our previous study (Jagannathan et al., 2024), we partitioned our data into three separate brain regions: central brain (inboard of the polarity reversal by 8 channels, typically around channel 3), middle (inboard by 3, typically around channel 8) and periphery (outboard by 3, typically around channel 14).

**Figure 1.**
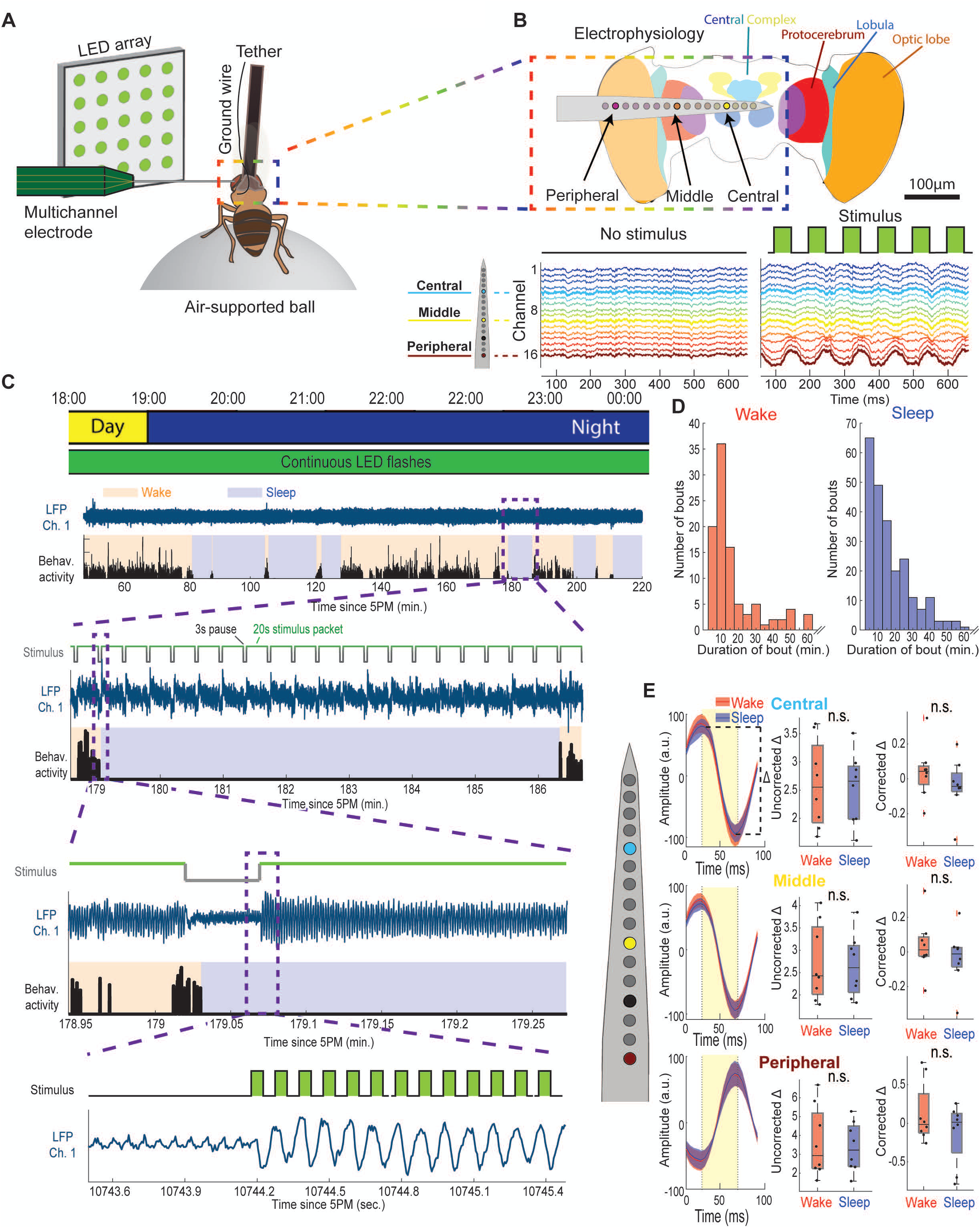
Visually-evoked local field potential (LFP) activity across the fly brain during wake and sleep. **A.** Schema of a tethered fly positioned on an air-supported ball. An LED array projects visual stimuli to the left eye, which also has a 16-channel multi-electrode inserted. **B.** Top, Schematized fly brain, showing the approximate position of electrode channels neuroanatomically. Marked are exemplar Central, Middle, and Peripheral channels, showing their typical positions during a recording. Bottom, Example LFP activity from an individual fly during an unstimulated recording period (left) and a period with a 10Hz visual stimulus (right). **C.** Top, Schematized representation of recording occurring across a day/night cycle. Underneath, the presence of stimuli across time is represented. Second from top, Example data from an early portion of an individual recording. Shown is LFP activity from an exemplar channel (blue), as well as fly movement activity (black). Shading indicates periods when the fly was determined to be waking (orange) or sleeping (blue), based on the movement activity. Third from top, Subset of the above, from a span of around 7 minutes, showing the timecourse of stimulus blocks (top, grey/green trace, where green represents periods of stimulus delivery), LFP activity from an example channel (middle, blue), and movement activity (bottom, black). Fourth from top, Zoom of the above activity, showing a transition from wake to sleep, as well as the onset of a stimulus bout. Bottom, A two second period of the above, showing stimulus-evoked activity. In a central channel. The top trace (‘Stimulus’) represents stimulus event onset and offset timing. **D.** Distribution of bout durations for Wake (left, red), and Sleep (right, blue) (N=8 flies). **E.** Left column, Averaged LFP (N=8) in response to visual stimulus presentation during wake (red) and sleep (blue). Stimulus timecourse is displayed with yellow shading. Top, middle, and bottom respectively relate to LFP activity in Central, Middle, and Peripheral channels. Middle column, Delta amplitude between peak and trough of average LFP during wake and sleep (n.s., One-way ANOVA with Bonferroni correction; N=8). Dots represent individual fly data. Right column, Mean-corrected delta amplitude of LFPs during wake and sleep (n.s., One-way ANOVA with Bonferroni correction; N=8).

Brain recordings were performed over several hours, spanning the natural transition to night-time sleep (Jagannathan et al., 2024, see Methods). Visual stimuli consisted of an 8x8 array of green light emitting diodes (LEDs) positioned 16 cm in front of the fly, slightly to the left and thus ipsilateral to the recorded brain hemisphere (Figure 1A). Visual stimuli were presented throughout the experiment at 10Hz (50ms duty cycle), in packets of 20s alternating with 3s pauses (Figure 1C). The stimuli produced robust steady-state visually-evoked potentials (SSVEPs) throughout the fly brain (responses in the central brain are shown in Figure 1C, bottom panel). We examined the behavioral data for evidence of sleep, defining sleep as periods of behavioral quiescence lasting 5min or more (see Methods).

Remarkably, flies were able to sleep while presented with this continuously flickering visual stimulus, displaying median sleep bout durations of 12.8 min and wake bout durations of 11.2 min (Figure 1D; an example sleep bout flanked by waking activity is shown in Figure 1C, second panel from the top). However, we found no significant difference in the average SSVEP amplitude during sleep compared to wake, for any of the three broadly-defined brain regions (Figure 1E), even after correcting for individual fly SSVEP amplitude variability (Figure 1E, right column; see Methods). This suggests that behavioral state does not modulate the average LFP response to the repetitive visual stimuli in this paradigm.

### Decreased responsiveness to deviant stimuli

We next questioned whether the probability of a stimulus occurring might affect the response across the fly brain. In the preceding experiment, the probability of a 50ms green flash occurring within the 20s visual sessions was 100%. To test whether stimulus probability altered visual responsiveness, we introduced a lower probability ‘deviant’ stimulus of a different color within the train of 10Hz stimuli (Figure 2A). Thus, a blue deviant stimulus was presented after every four green stimuli, such that the green ‘carrier’ stimuli still occurred at 10Hz while the blue deviant stimuli occurred at 2Hz. The experiment was counterbalanced so that for some 20s sessions the blue stimuli were deviant, while in other sessions the green stimuli were deviant. This arrangement allowed us to compare the responses to carriers and deviants strictly within a color, to potentially account for any differences in color processing. Thus, we asked whether the LFP response to a green or blue deviant occurring at 2Hz was different from the response to a green or blue carrier occurring at 10Hz. To ensure that our comparisons were historically identical and only contingent on the stimulus probability, we specifically compared carriers preceded by a deviant to (identically colored) deviants preceded by a carrier (because both conditions involved a color change). We first only examined responses during waking epochs, to determine if responses varied in an aroused brain. An example of LFP responses across the brain to carrier and deviant stimuli is shown in Figure 2B. We found that stimulus probability had a significant effect on SSVEP amplitude: in the central brain, responses to deviant stimuli were smaller than responses to carrier stimuli of the same color, for both colors (Figure 2C, top row). Interestingly, this effect was lost in middle and peripheral channels for blue deviants but robustly maintained for green deviants (Figure 2C, middle and bottom rows). Blue and green are show some differences arising at the level of photoreceptors that project to distinct medulla neurons in the fly optic lobes (Yamaguchi et al., 2010), potentially giving rise to different SSVEP dynamics.

**Figure 2.**
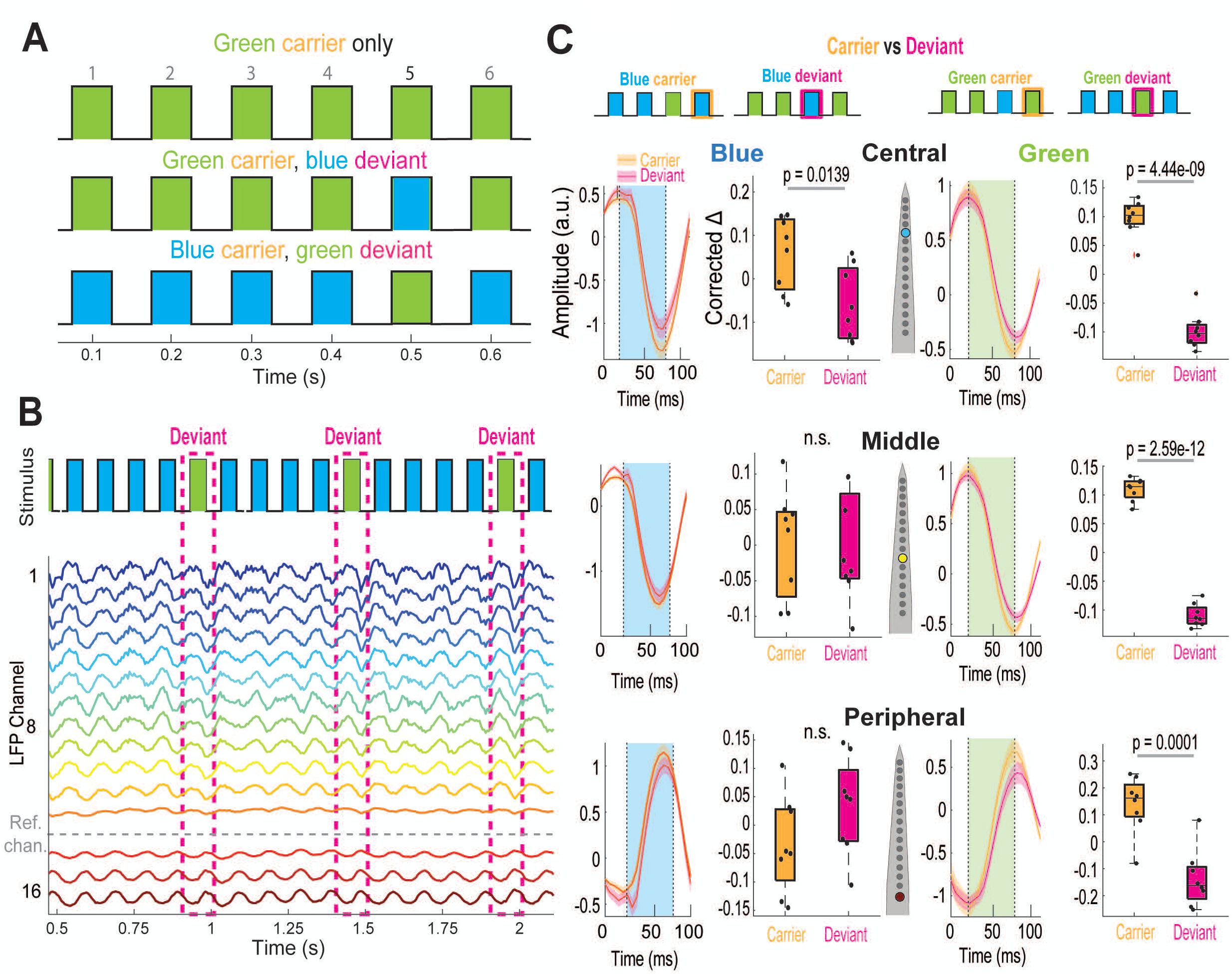
Decreased responsiveness to deviant stimuli. **A.** Schematized stimulus blocks, showing periods with a continuously repeated carrier (top), and those with color deviants (middle and bottom). Numbers above schema represent position in sequence of events. All conditions are counterbalanced for color across individuals. **B.** Stimuli and corresponding LFP activity across the fly brain during a portion of a stimulus block, showing stimulus event identity and matching LFP. Deviant events are highlighted with a dashed magenta box. Grey dashed line indicates the polarity reversal channel for this fly, which is used for as a reference channel for noise subtraction. **C.** Left, LFPs and corrected delta amplitudes across channel in response to a blue carrier (orange shading) or blue deviant (magenta shading). Right, as with left, but for green carriers/deviants. P-values where indicated represent one-way ANVOA with Bonferroni correction (N=8).

### The central brain is insensitive to carrier effects

Not all carrier stimuli within a color are equal in the context of our paradigm. While their luminosity and hue are identical, their history is different (Figure 2A,B, top row schema). Thus, carriers #2-4 are all preceded by another carrier of the same color, while carrier #1 is preceded by a differently-colored deviant. In this regard, carrier #1 is also deviant-like (a nature we exploited for appropriate comparison with the deviant, earlier). Interestingly, we found that carrier #1 evoked a significantly different response than carriers #2-4 in the optic lobe (Figure 3A,B, bottom panels). Colors had opposite effects: blue carrier #1 was significantly smaller than the subsequent 3 blue carriers, while green carrier #1 was significantly larger than the 3 subsequent green carriers. This suggests a color change effect, which might be expected considering distinct color-processing pathways in the optic lobes of the *Drosophila* visual system (Paulk et al., 2013). This effect persisted in the middle channels, although it was only significant for green carriers (Figure 3A,B, middle panels). Interestingly, the carrier effect was lost in the central channels (Figure 3A,B, top panels). This suggests that SSVEPs recorded in the central fly brain are tracking stimulus probability (Figure 2C) rather than changes in stimulus color.

**Figure 3.**
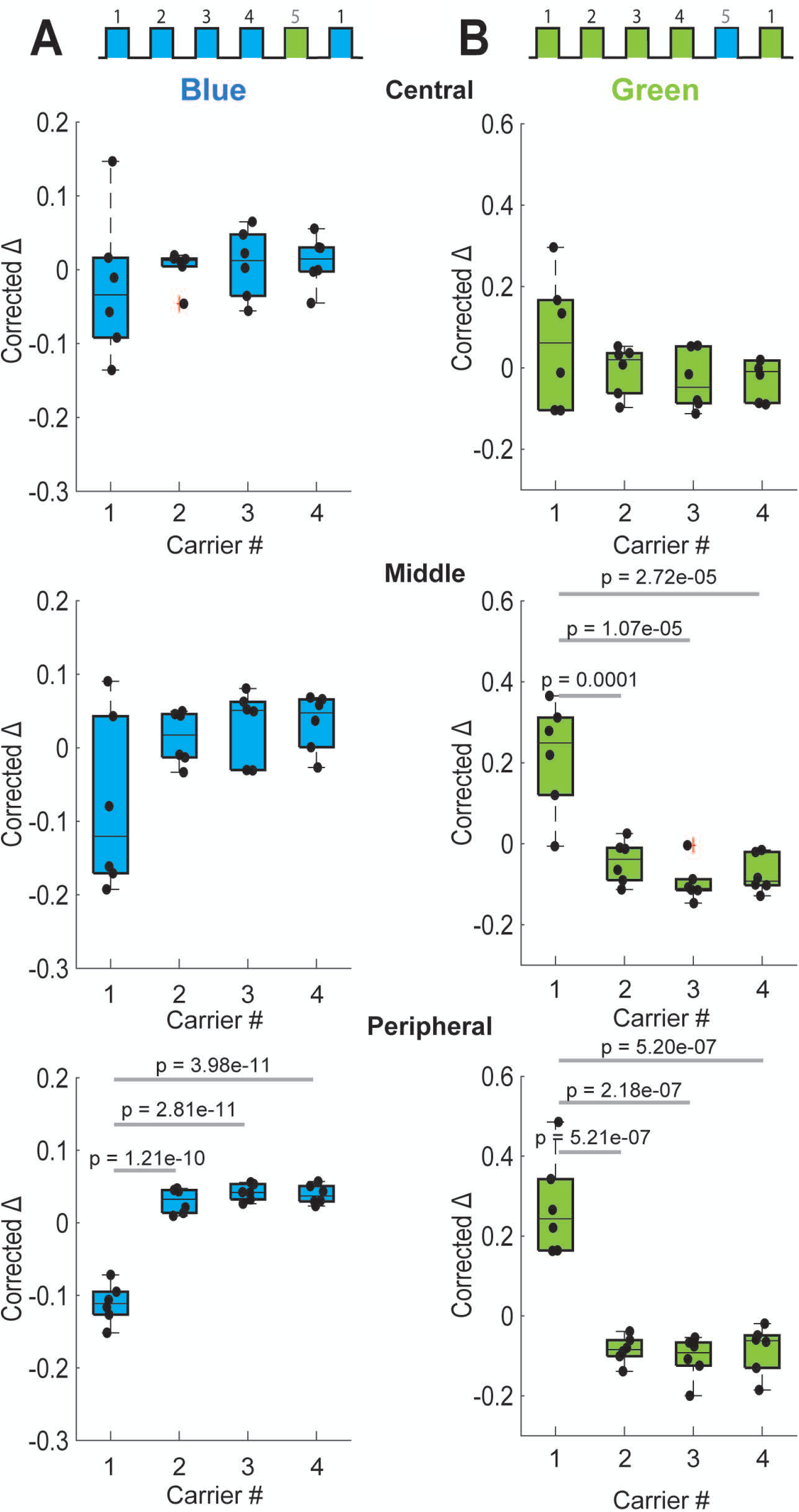
The central brain is insensitive to carrier effects. **A.** Comparison of corrected delta amplitude for the first through to fourth blue carrier after a green deviant, across channels. Significance between groups is indicated with grey bar and a p-value, based on one-way ANOVA with Bonferroni correction (N=8). **B.** As with A, for green carriers after a blue deviant.

### Stimulus probability rather than uncertainty determines SSVEP response amplitude

In the preceding paradigm, both the deviant and carrier stimuli are equally predictable; the deviant occurs after exactly every four carriers. We next questioned whether decreasing the predictability of the deviant might alter how the fly brain responds to it. To test this, we jittered the position of the differently colored deviant stimulus such that it could appear after different numbers of carrier stimuli (Figure 4A, left panel), while on average still occurring at 2Hz compared to the 10Hz carrier (Figure 4A, right panel). All experiments were counterbalanced like before so that comparisons could be made within a color (i.e., green jittering deviant compared to green carrier, blue jittering deviant compared to a blue carrier). We found that a jittering deviant also evoked smaller responses compared to carrier SSVEP amplitudes (Figure 4B, left panel), and the combined data was significant across the fly brain (Figure 4B, right 3 panels). To further investigate whether predictability affected SSVEP amplitudes, we specifically examined only the jittering the subset of deviants that happened to occur after exactly 4 carriers (Figure 4C, left panels), thus providing a physically identical comparison for the non-jittering experiments presented in Figure 2. A spectral analysis of the visual stimulus shows how different the context is for these two different paradigms (Figure 4C, right panels). Nevertheless, when restricted to comparing identical physical histories, we found no significant difference between a jittering and a non-jittering deviant (Figure 4D). This suggests that probability rather than predictability is driving the deviant effect in this particular paradigm.

**Figure 4.**
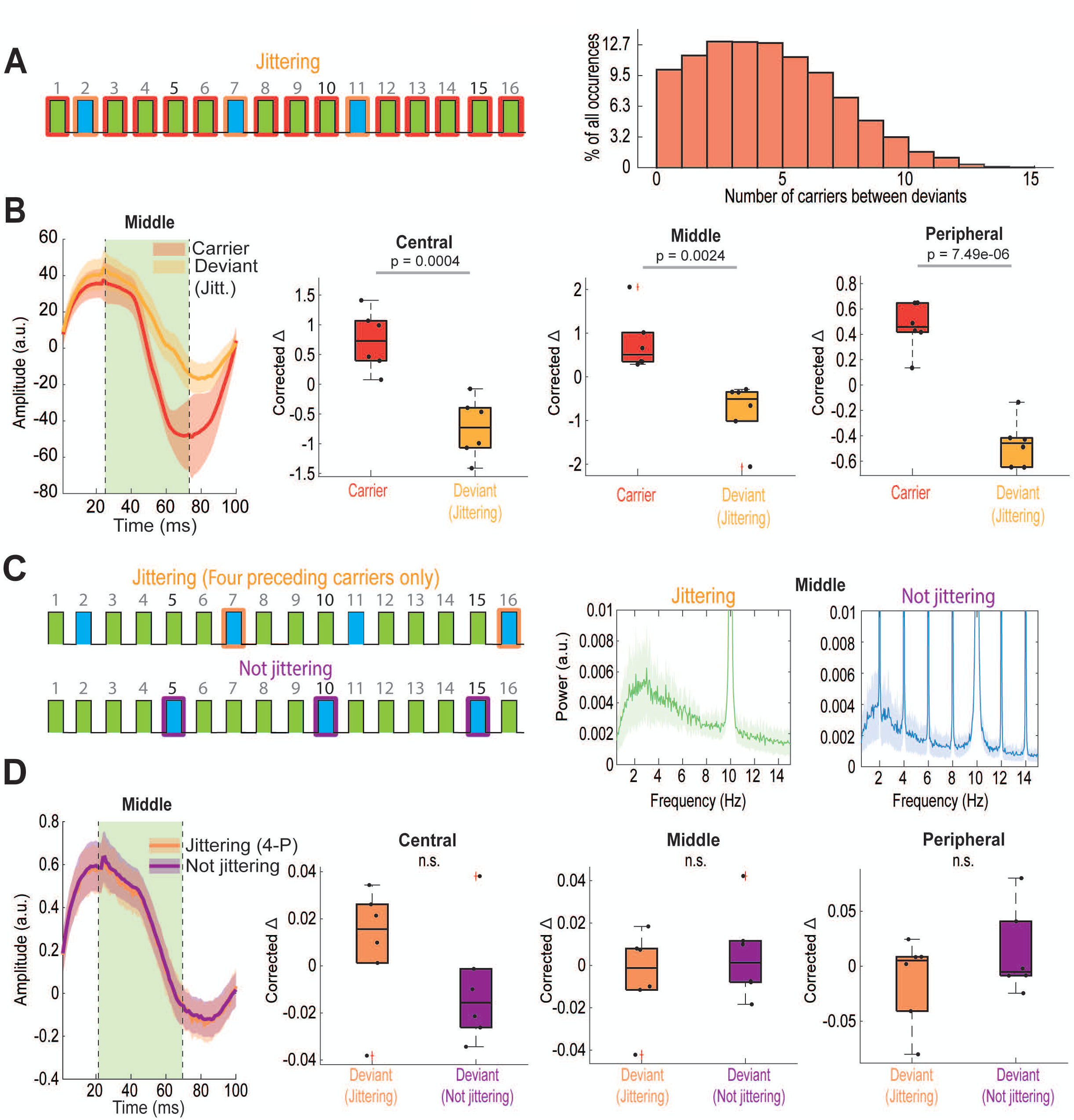
Stimulus probability rather than uncertainty determines LFP response amplitude. **A.** Left, Example stimulus schema for a block where the position of the deviant in sequence is ‘jittered’ according to a normal distribution. Orange boxes indicate example deviant positions, while red boxes indicate carriers used for comparison. Right, calculated actual distribution of deviants in sequence (N=6). **B.** Left, Average LFPs across individuals for carriers of any position (red) versus deviants jittered in sequence (yellow). Colors are combined here, such that carriers could either have been green or blue, and similar for deviants. Middle through Right, Comparison of corrected delta amplitude values for carrier and deviant conditions mentioned above, across Central, Middle, and Peripheral channels. P-values indicated on plot represent one-way ANOVA with Bonferroni correction. Dots each represent an individual fly average (N=6). **C.** Left, Example schematized stimulus sequences for a ‘jittering’ and ‘not jittering’ block. Colored boxes respectively indicate the selection of a deviant stimulus event that happened to be preceded by 4 carrier events (orange), as well as a non-jittering deviant that would always be preceded by 4 carrier events (purple). Right, Power spectrum of LFP activity in middle channels during jittering and not jittering blocks, normalized and averaged across individuals. F1 carrier frequency can be observed at 10Hz, while F2 deviant frequency peaks at 2Hz. **D.** As with B, but only comparing jittering (orange) and non-jittering (purple) deviants both preceded by 4 carrier events (one-way ANOVA with Bonferroni correction; N=6).

### Spontaneous sleep decreases the brain response to deviants specifically

In our earlier experiments, we found that spontaneous sleep had no effect on the amplitude of the SSVEP response to 10Hz carrier stimuli presented throughout the night (Figure 1). Knowing that the waking fly brain’s response to deviant stimuli is different than its response to carriers, we next examined whether these were processed differently during spontaneous sleep compared to wake. Since we observed the same result with deviants irrespective of whether they were jittering or predictable, we focused on our original design where deviants occurred at exactly 2Hz (non-jittering) within a 10Hz carrier of a different color. Flies were exposed to these stimuli during long-term multichannel recordings as before (these were the same overnight flies as in Figure 1, which were also exposed to 20s packets of only carrier stimuli), and sleep was identified as periods of quiescence lasting 5min or more under these conditions (Figure 5A). As before, we only compared effects within a color, thus green deviants to green carrier #1). We found that sleep significantly decreased the SSVEP response to deviant stimuli, while (consistent with Figure 1 results) responses to carrier stimuli were still not significantly affected by behavioral state (Figure 5B). Interestingly, the effect of sleep on deviant responses was only significant in the central and middle brain channels and was not significant in the peripheral (outer optic lobe) channels. Together, this suggests that processing of high-probability stimuli (i.e., carriers) are unaffected by sleep throughout the fly brain, while processing of low probability (deviant) stimuli is affected by sleep in the more central regions of the fly brain.

**Figure 5.**
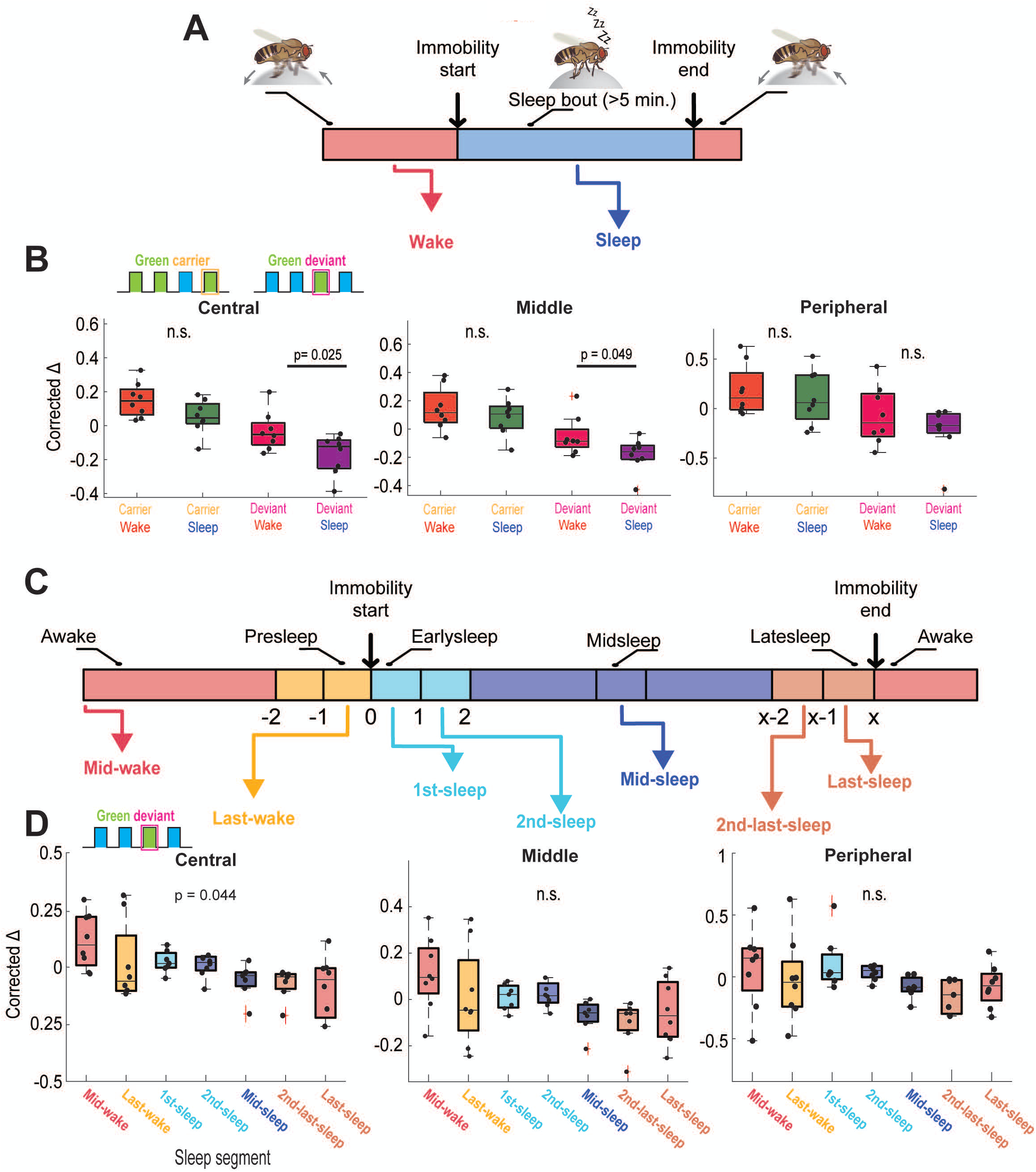
Spontaneous sleep decreases the brain response to deviants specifically. **A.** Schema illustrating the conversion of locomotive activity data into wake and sleep bout identification. **B.** Corrected delta amplitude values for green carrier and deviant responses during wake and sleep. Comparisons are conducted only within group (e.g. Carrier Wake vs Carrier Sleep) and represent one-way ANOVA with Bonferroni correction (N=8). **C.** Schema separating wake and sleep bouts into segments based on time since sleep onset (Jagannathan et al., 2024; Yap et al., 2017). Each segment represents a particular time period either before/after the onset of immobility or during sleep. **D.** Corrected delta amplitude values for green deviant stimulus responses during the time segments listed in C. P-values listed on plot represent the overall significance of one-way ANOVA with Bonferroni correction (N=8).

### Spontaneous sleep stages have different effects on the response to deviant visual stimuli

In a recent study using the same multichannel brain recording preparation, we identified distinct stages of sleep based on classification of electrophysiological activity as well as on microbehaviors (Jagannathan et al., 2024). We therefore questioned whether the attenuated response to deviant stimuli was characteristic of any specific phase of sleep. To address this, we partitioned all identified sleep epochs (>5min) into five segments (Figure 5C): the first minute of sleep (1^st^ sleep), the second minute of sleep (2^nd^ sleep), the middle minute of sleep (mid-sleep; variable in position across bouts of different durations), the second-last minute of sleep (2^nd^-last-sleep), and the last minute before awakening (last sleep). The randomized distribution of our visual paradigm throughout the night ensured that all counterbalanced blue/green presentations occurred multiple times for each sleep segment, for each fly (see Methods). For comparison, we performed the same analyses for two wake segments: data taken from the middle minute of a preceding wake epoch (mid-wake), and data taken from the last minute of preceding wakefulness (last-wake), when flies were still moving (Figure 5C). We found that only recordings taken from the central brain revealed significant differences among this segmented data (Figure 5D), however with no specific sleep epoch significantly different on its own compared to other epochs.

In the preceding analyses, sleep stages are inferred by their timing following the beginning of a spontaneous sleep bout. While this has been indicative of when active or quiet sleep stages might occur (Yap et al., 2017), we sought an alternative behavioral approach to disambiguate distinct sleep stages in our data. In a recent study, we found that flies displayed more rhythmic proboscis extensions during their deep sleep stage (Jagannathan et al., 2024), which supports an earlier finding (van Alphen et al., 2021). We therefore wondered whether responses were affected during the deep sleep epochs characterized by bursts of rhythmic PEs. Since rhythmic PEs also occur during wake in this preparation (Jagannathan et al., 2024), this provided an ideal comparison to determine whether behavioral state (i.e., deep sleep) or just the behavior itself (i.e., PEs) governed the weakened response to deviant visual stimuli. As previously (Jagannathan et al., 2024), we employed machine learning approaches to identify PEs in our overnight recordings of flies exposed to visual stimuli, while also tracking leg movements to monitor sleep (Figure 6A, top). We identified all PE activity that qualified as rhythmic and aligned them in time with our multichannel electrophysiological activity (Figure 6A, bottom). Closer examination of proboscis activity revealed two kinds of movements, either rhythmic (proboscis extensions, PEs) or individual, questing-like ‘reaches’, which primarily happened during wake (Figure 6B, top panels). Given their purported relevance for sleep functions (van Alphen et al., 2021) we only examined PEs. Since PEs themselves affect fly brain activity (Jagannathan et al., 2024), we only examined SSVEPs that were in between the rhythmic PE events (Figure 6B, bottom panels), to determine if the PE ‘context’ (during wake vs sleep) affected SSVEP amplitude (see Methods). We identified rhythmic PE events in all of our data, allowing us to partition these into the same temporal segments as outlined above. Consistent with our results so far, sleep had no effect at all on how carrier stimuli were processed (Figure 6C). In contrast, deviant stimuli were sensitive to sleep quality. Interestingly, only PEs occurring during mid-sleep epochs (when flies are in deep sleep; Jagannathan et al., 2024; van Alphen et al., 2021) were associated with significantly decreased responsiveness to deviant stimuli in the central brain, although other sleep epochs showed a trend in the same direction (Figure 6D). This effect was not seen in the optic lobe recordings, while data taken from middle channels seemed intermediate. This suggests that during the deepest phase of sleep, when flies typically display the most rhythmic bursts of PEs (Jagannathan et al., 2024), the central fly brain is significantly less responsive to low-probability stimuli but equally responsive to high probability stimuli.

**Figure 6.**
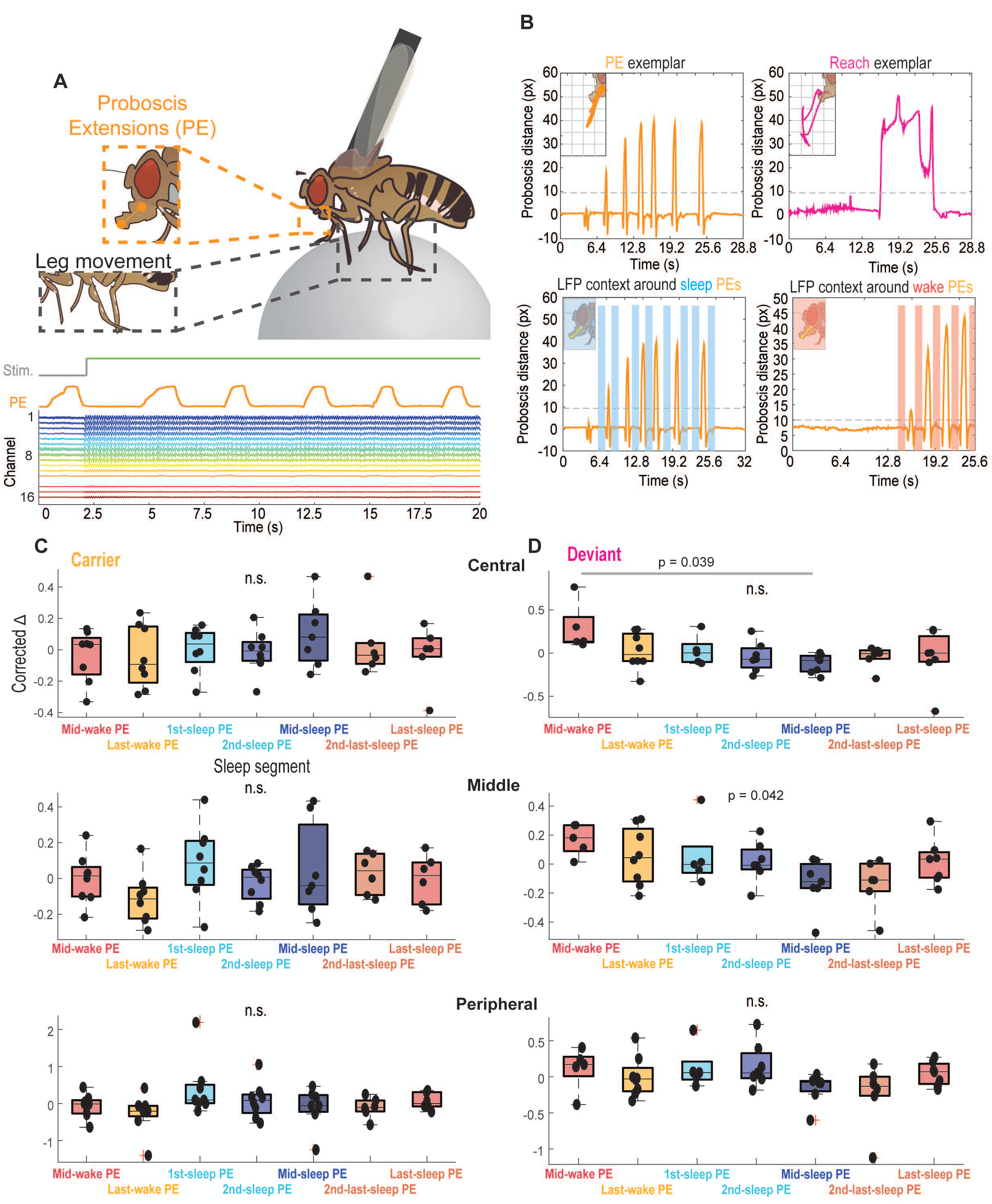
Decreased response to deviant stimuli during deep sleep specifically. **A.** Top, Schema of a tethered fly, focusing on the tracking of proboscis extensions (PE, orange), and locomotive activity via leg movement (black). Bottom, LFP activity (colored traces) in an example fly, during a spell of proboscis extensions (orange trace). Minimal PE artifact is evident in the brain recordings. The presence of visual stimuli during this period is indicated above (grey/green trace). **B.** Top Left, Exemplar proboscis extension distance tracked from resting position (orange) during a PE spell. Inset, a trace of the position of the proboscis in 2D space during the event. Top Right, As with Top Left, for a ‘reach’ event, showcasing the longer time course and singular nature. Bottom Left, Exemplar PE spell during a sleep bout, with inclusion of shading to show portions of time when LFP activity was sampled. Bottom Right, As with Bottom Left, for a PE spell occurring during a wake bout. **C.** Corrected delta amplitude values for green carrier events occurring during PEs of specific sleep segments, across different brain regions. Dots represent individuals (N=8). Significance calculated via one-way ANOVA with Bonferroni correction. **D.** As with C, for green deviant events. Significant comparisons between groups represented with grey shading while overall significance presented purely as text (one-way ANOVA with Bonferroni correction; N=8).

## DISCUSSION

Convincing evidence that Drosophila flies really sleep was first provided using mechanical stimuli, which showed that flies had increased arousal thresholds to these stimuli if they had been inactive for 5 minutes or more (Shaw et al., 2000). Subsequent experiments using mechanical stimuli confirmed 5 minutes immobility as a useful threshold for studying sleep in this model (Huber et al., 2004; van Alphen et al., 2013), although recent studies have questioned whether sleep quality is adequately addressed by this proxy (Chowdhury et al., 2023). Other recent studies have also used olfactory stimuli to assess sleep quality and intensity in Drosophila (French et al., 2021). Here, we used visual stimuli to investigate sensory processing during sleep, and we focused on LFP responses across the fly brain rather than behavior. We found that repetitive visual stimuli are processed similarly during sleep and wake, but lowering the probability of a repetitive stimulus made it less detectable in the central brain during sleep. This effect reached a nadir during the later stages of sleep, when flies displayed rhythmic proboscis extensions. This confirms for another modality (vision) that flies are probably sleeping after 5 minutes of immobility. This also has relevance for uncovering sleep functions: deviants are typically surprising, and one proposed function of sleep is to optimize how prediction errors are processed (Van De Poll and van Swinderen, 2021). It is therefore possible that altered processing of deviant stimuli during sleep also accomplishes a homeostatic function. Interestingly, the deviant effect was only evident in the central brain recordings, even though the entire fly brain – even the optic lobes – was found to display distinct sleep features in multichannel LFP recordings (Jagannathan et al., 2024). This suggests that any mechanism suppressing the detection of deviant visual stimuli likely resides in the central brain, while ‘sleeping’ optic lobes are still able to process these deviant stimuli in the same way as during waking. This is consistent with previous studies uncovering distinct sleep-related features the central brain activity (Yap et al., 2017).

One surprising finding from our study was that deviant visual stimuli evoked smaller LFP responses in the fly brain, compared to high-probability carrier stimuli. In mammalian studies, deviant stimuli typically evoke larger responses in LFP or EEG recordings (Shiramatsu and Takahashi, 2021). Indeed, such amplified EEG effects have formed a foundation for studying predictive processing in humans and other mammals, whereby unexpected events evoke a comparatively larger ‘dip’ in brain recordings, termed ‘mismatch negativity’ (MNN; Näätänen, 1990). In the fly brain, we found nothing resembling MNN. Rather, we found the opposite: a decreased response even to a properly unpredictable stimulus, and a further attenuated response to deviant stimuli during sleep – but only in the central brain. How might this discrepancy with similar readouts in other animals be explained? One explanation lies in our unique recording preparation. In our multichannel recording setup, the linear array of 16 recording sites are oriented along a specific plane, facing the back of the fly’s head (see Figure 1B). Field potential electrodes only record what they are allowed to detect, namely parallel electrical fields lying in a specific orientation so as to reveal a current and thus a voltage differential. It is therefore possible that other electrical fields orthogonal to the detected field are increasing in amplitude with certain stimuli (e.g., deviants) and thereby cancelling to some extent the fields we are measuring in the current experimental design. This is a known drawback of electrophysiology: orthogonal fields cancel out, and even an LFP flatline does not necessarily indicate a lack of neural activity (Buzsáki et al., 2012). In our study we have detected an effect of deviant stimuli, which necessarily means distinct processing for those stimuli – irrespective of how that processing is manifested in LFP activity. Future studies, for example using sharp recordings in the central brain rather than linear probes across the brain, should reveal whether deviant visual stimuli always evoke smaller LFP responses in Drosophila, or whether this might indeed just be a feature of the orientation of the multichannel recording electrodes used in this study.

Another surprising finding from our study was that making the deviant less predictable did not alter the LFP response, compared to a predictable deviant with the same historical features. One might have expected an unpredictable deviant to be more surprising, therefore evoking an even more attenuated version of the deviant response. Rather, if anything, there was trend to a larger response in the jittering deviant, in the central brain (Figure 4D). This again calls for employing a variety of approaches for studying predictive processing in the central fly brain, now that we have shown that identical visual stimuli are not all processed equally depending on behavioral state and history. Alternate LFP recording orientations (Grabowska et al., 2020; van Swinderen and Greenspan, 2003) as well as calcium imaging (Tainton-Heap et al., 2021; Troup et al., 2023), coupled with transient manipulations of visual processing circuits in the central brain, should further elucidate the nature of the of the deviant effects we have uncovered in this study.

## ACKNOWLEDGEMENTS

The authors thank Travis Jeans for help with the multichannel recording preparation, Sridhar Jagannathan for help with behavioral data analysis, Leonie Kirszenblat for comments on the manuscript, and members of the van Swinderen laboratory for feedback on the work.

## DATA AND RESOURCE AVAILABILITY

These data will be made available upon publication in a searchable database.

